# TAF15 amyloids propagate via defined motifs in a prion-like fashion

**DOI:** 10.1101/2025.11.17.688886

**Authors:** Katerina Konstantoulea, Laxmikant Gadhe, Frank Goodavish, Ankit Gupta, Harichandra D. Tagad, Jaime Vaquer-Alicea, Nikhil B. Ghayal, Shanu F. Roemer, Michael A. DeTure, Alissa L. Nana, Salvatore E. Spina, Lea T. Grinberg, William M. Seeley, Dennis W. Dickson, Charles L. White, Marc I. Diamond, Nikolaos N. Louros

## Abstract

TATA-box binding protein–associated factor 15 (TAF15) is an RNA-binding member of the FET family recently identified as the primary fibrillar constituent in a subset of frontotemporal lobar degeneration (FTLD-FET) cases. Although TAF15 is also linked to amyotrophic lateral sclerosis (ALS), the molecular basis and propagation behavior of its aggregates remain unknown. In this work, we show that recombinant TAF15 forms amyloid fibrils under physiological conditions and developed a single-fluorophore TAF15 biosensor to quantitatively monitor their cellular propagation. Using this system, we demonstrate that both recombinant TAF15 fibrils and pathological aggregates extracted from atypical FTLD with ubiquitin inclusions (aFTLD-U) patient brains seed aggregation efficiently and transmit serially between cells, demonstrating hallmark features of prion-like propagation. Seeding was specific to TAF15 and absent for other amyloidogenic proteins, including the homologous protein fused-in-sarcoma (FUS), revealing an unexpected cross-seeding barrier. Occasional colocalization of FUS within TAF15 inclusions was observed upon transient co-expression, suggesting that FUS can be passively recruited rather than acting as an inducer of pathology in FTLD-FET brains. Computational and peptide-based experimental mapping identified multiple aggregation-prone regions within the TAF15 low-complexity domain that coincide with hotspots stabilizing the core of ex vivo TAF15 amyloid fibrils. These short motifs encode the propagation propensity of TAF15 aggregation in vitro and in cells. Together, these findings establish TAF15 as a bona fide amyloid-forming, prion-like protein and define the sequence grammar underlying its self-assembly, providing a mechanistic framework for its role in FTLD-FET and ALS and offering tractable molecular targets for therapeutic intervention.

## Introduction

Frontotemporal lobar degeneration (FTLD) is a leading cause of early-onset dementia^1^, characterized by progressive deterioration of behavior, language, and executive function, often overlapping with motor neuron diseases such as amyotrophic lateral sclerosis (ALS)^2-4^. The disorder imposes a profound burden on patients and caregivers, and its molecular diversity complicates diagnosis and therapy^5^. Overall, FTLD is pathologically classified into three main subtypes: FTLD-tau, FTLD-TDP, and FTLD-FET, based on the predominant protein found in neuronal inclusions. FTLD-FET, which accounts for approximately 10–20% of cases, is defined by cytoplasmic inclusions composed of FET family proteins: FUS, EWSR1, and TAF15^6-9^.

Historically, FUS was considered the principal aggregating species and driver of pathology in FTLD-FET^10-12^, a view influenced by the discovery that FUS mutations can cause ALS in the absence of frontotemporal degeneration^13,14^, and by early reports showing that recombinant FUS can assemble into amyloid-like fibrils in vitro^15^. However, no FUS mutations have been associated with FTLD to date^16^, and FUS filaments have never been isolated from patient brain tissue. In contrast, recent cryo-electron microscopy (cryo-EM) studies revealed that TAF15, not FUS, constitutes the major fibrillar component in the frontal and temporal cortices of affected individuals^17^, further prompting the re-classification of many cases previously diagnosed as FTLD-FUS into FTLD-FET. Consistent with this, inclusions in FTLD-FET are immunoreactive for TAF15 and transportin-1, and in a subset of cases also for EWSR1^6-9,18^. The predominant subset of FTLD-FET, referred to as atypical FTLD with ubiquitin-positive inclusions (aFTLD-U), is not related to FUS mutations and is characterized by TAF15-positive inclusions that predominantly affect the frontal and temporal cortices^16^. Notably, in contrast to FUS, TAF15 aggregates uniquely correlate with disease severity and cognitive decline in aFTLD-U cases^17^. These findings suggest that TAF15 is not a secondary inclusion marker^7^ but a central molecular driver of FTLD-FET pathogenesis.

TAF15 is a multifunctional RNA-binding protein that shares domain architecture and sequence composition with its FET homologs^19^. Like FUS and EWSR1, it contains a glycine- and tyrosine-rich low-complexity domain (LCD) capable of phase separation and stress granule assembly, followed by an RNA recognition motif and C-terminal zinc finger^20,21^. Both wild-type and mutant forms of TAF15 have been implicated in cytoplasmic inclusion formation in ALS^22-28^. ALS-linked variants, including A31T, N49S, S51G, E71G, and M92V, enhance aggregation in cellular and animal models^22,29^, while knockdown of TAF15 or parkin-mediated degradation mitigates motor deficits and neurotoxicity^24^. Inclusions immunoreactive for TAF15 are also observed in related conditions such as neuronal intermediate filament inclusion disease (NIFID-FET) and basophilic inclusion body disease (BIBD)^7-9,30,31^. Together, these findings identify TAF15 as a mechanistically underexplored yet increasingly relevant contributor to neurodegenerative proteinopathies.

Despite these advances, the molecular determinants that drive TAF15 aggregation and its potential for prion-like propagation remain unknown. In other amyloid-driven disorders, short aggregation-prone regions (APRs) act as critical nucleation elements that define fibril architecture and polymorphism^32-39^. Such β-strand-forming motifs stabilize the amyloid core^40,41^ and are sufficient to induce aggregation when isolated^36-38,42^. Proteins such as tau, TDP-43, and α-synuclein harbor distinct APRs that dictate strain behavior and propagation dynamics^32^, as well as therapeutic tractability^43-45^. Whether TAF15 contains similar sequence-encoded motifs that endow it with self-propagating properties has remained completely unaddressed.

Here, we define the molecular and cellular mechanisms underpinning TAF15 aggregation and propagation. Using biophysical, computational, and cellular approaches, we show that recombinant TAF15 forms amyloid fibrils under physiological conditions and that pathological aggregates from aFTLD-U patient brains seed aggregation efficiently and serially in cells, establishing that TAF15 propagates via a prion-like mechanism. To quantify and visualize these events, we developed a single-fluorophore mEOS3.2-based TAF15 biosensor, which reports aggregation with high sensitivity and specificity, unresponsive to other amyloidogenic proteins, including the homologous FUS. Thermodynamic and peptide-based mapping identified several discrete APRs within the TAF15 LCD that coincide with stabilizing hotspots in patient-derived fibril structures and act as nucleation elements both in vitro and in cells. Together, these findings establish TAF15 as an amyloid-forming, prion-like protein and reveal the sequence determinants that underlie its selective vulnerability in FTLD-FET and ALS.

## Results

### In vitro formation of TAF15 amyloid fibrils

Previous structural analysis of FTLD-FET brain tissue identified residues 7–99 of TAF15 as forming the amyloid core of the fibril deposits^17^. To determine whether this region is sufficient to drive amyloid assembly in vitro, we expressed the TAF15(7–99) fragment recombinantly using affinity purification (**Fig. 1A**). After tag removal, SDS–PAGE analysis showed a single band corresponding to the cleaved protein (**Fig. 1B**). When incubated under physiological buffer conditions (100µM, PBS, 37 °C), the TAF15 fragment spontaneously assembled into uniform, unbranched amyloid fibrils visualized by transmission electron microscopy (TEM) (**Fig. 1C**). Thioflavin-T (ThT) endpoint fluorescence assays confirmed the propensity of the fragment to self-assemble into amyloid fibrils (**Fig. 1D**).

**Figure 1.**
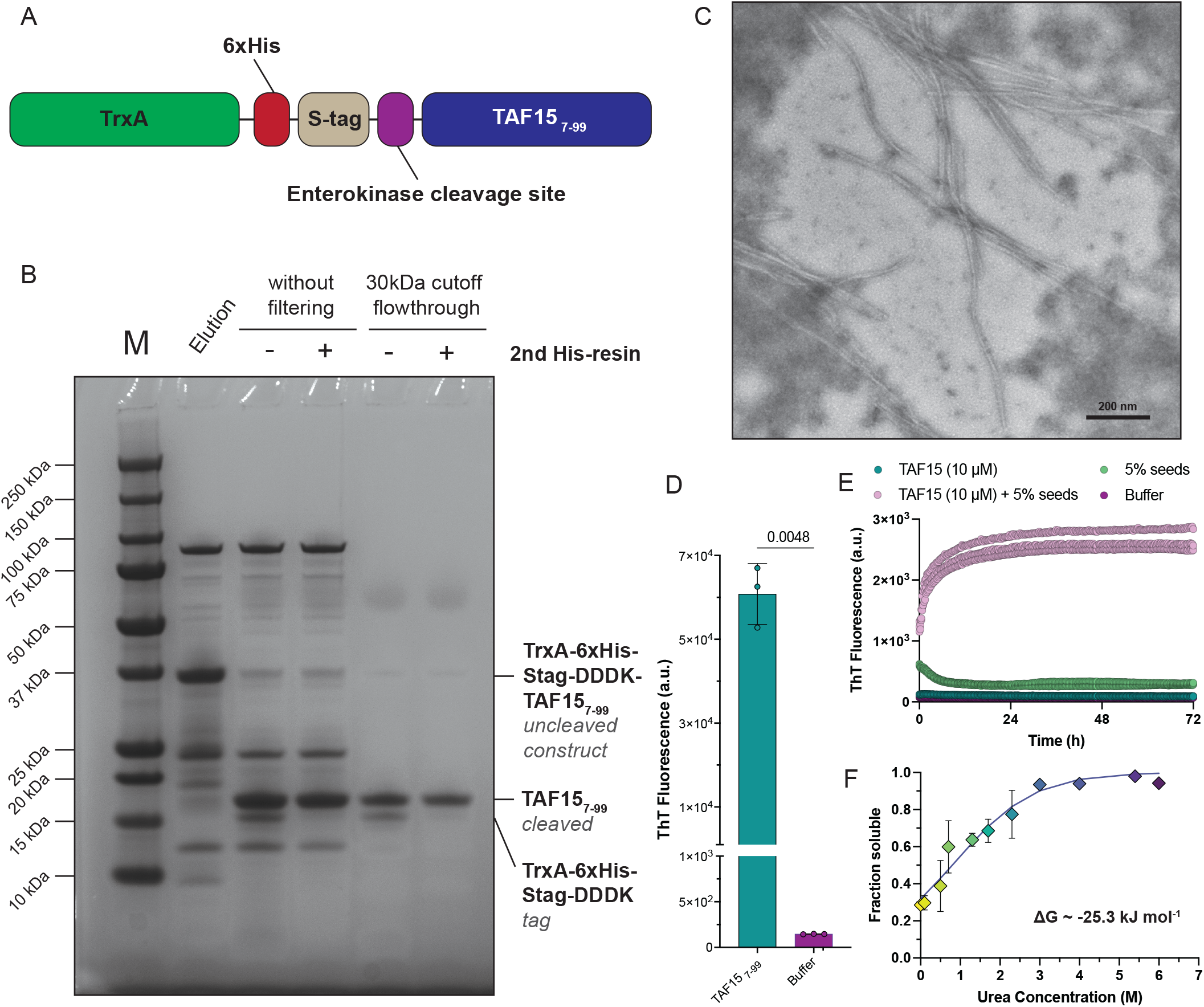
Recombinant TAF15 forms amyloid fibrils under physiological conditions. **(A)** Schematic of the recombinant TAF15 construct (residues 7–99) used for production and aggregation assays. **(B)** SDS–PAGE of purified TAF15(7–99) before and after tag cleavage showing a single band at the expected molecular weight. **(C)** Negative-stain TEM of end-state assemblies reveals unbranched, fibrillar morphologies typical of amyloid fibrils. Scale bar: 200nm. **(D)** Endpoint ThT fluorescence of TAF15(7–99) reactions confirms amyloid formation under the indicated conditions. Statistics: Two-tailed t-test with Welch’s correction (n=3 individual replicates). **(E)** ThT kinetic assays (n=3 individual replicates) at 10 µM show no detectable self-assembly over 72 h in unseeded reactions, whereas the addition of preformed TAF15 fibrils induces rapid sigmoidal growth (seeded). **(F)** Stability profiling of TAF15 fibrils by chemical depolymerization with urea and measured by flow-induced dispersion analysis, reports the fraction of solubilized monomer as a function of denaturant.

We next examined whether preformed TAF15(7–99) fibrils can seed the aggregation of monomeric protein. Unseeded reactions at a lower concentration of TAF15 (10µM) showed no measurable ThT increase over 72 h, whereas the addition of preformed fibrils resulted in rapid, sigmoidal kinetics characteristic of templated nucleation (**Fig. 1E**). Together, these results demonstrate that the TAF15(7–99) fragment forms amyloid fibrils with typical amyloid self-propagating properties.

To assess the structural integrity of the assembled TAF15 fibrils, we analyzed fibril stability using chemical denaturation and Taylor dispersion analysis. This approach exploits differences in diffusion behavior between soluble and fibrillar species under laminar flow, allowing their transient incomplete separation (TIS) and quantification of the soluble fraction. Under defined hydrodynamic conditions, monomers display classical Taylor dispersion behavior, whereas the mass transfer of fibrils is dominated by convection^46^. Chemical depolymerization followed by deconvolution therefore enables direct estimation of fibril stability from the extent of solubilized material. FIDA analysis produced well-defined elution profiles. Quantification of the soluble fraction via integration of the monomer AUC relative to the total area of the elution profiles reflected concentration-dependent chemical denaturation (**Fig. 1F**). A global nonlinear fit of all data points resulted in agreement to the predicted curve, yielding a ΔG value of −25.3 kJ·mol^−1^, which is comparable to previously reported stabilities of pathological α-synuclein variants^46,47^.

### A TAF15 biosensor captures self-propagating amyloid aggregation in cells

To establish a cellular model of TAF15 aggregation, we designed HEK293T biosensors expressing the low-complexity domain (LCD) of TAF15 (residues 1–200) fused to the photoconvertible fluorophore mEOS3.2 (**Fig. 2A**). Western blotting of cell lysates revealed a clear band corresponding to the fusion protein, which was absent in control HEK293T cell lysates (**Fig. 2B**). Exposure of biosensor cells to recombinant TAF15(7–99) fibrils induced a strong, concentration-dependent FRET signal detected by flow cytometry (calculated EC_50_ value ∼ 92 nM monomer equivalent) (**Fig. 2C, purple curve**). Lysates generated from these seeded cells efficiently induced aggregation in naïve biosensors, demonstrating serial transmission of the TAF15 conformer (**Fig. 2C, yellow curve**). Control cell lysates generated from unseeded TAF15-mEOS3.2 biosensor cells failed to trigger any inclusion formation in naïve cells (**Fig. 2C, green curve**). High-resolution confocal imaging revealed cytoplasmic inclusions displaying fibrillar substructure (**Fig. 2D**). By performing sarkosyl extraction, we also isolated TAF15 aggregates from the biosensor cells. Morphological analysis revealed the formation of amyloid fibrils with widths around 70-100 Å, similar to what was previously reported for ex vivo TAF15 fibrils^17^ (**Fig. 2E**). To further validate the aggregation properties of the intracellular inclusions, we also performed fluorescence recovery after photobleaching (FRAP) experiments. FRAP across several puncta demonstrated loss of fluorescence with no recovery (**Fig. 2F–H**), indicating that the inclusions represent immobile fibrillar aggregates. Collectively, these experiments show that the mEOS3.2-based TAF15 biosensor faithfully reports TAF15 nucleation-dependent aggregation and provides evidence of the prion-like cellular propagation propensity of TAF15 amyloid fibrils.

**Figure 2.**
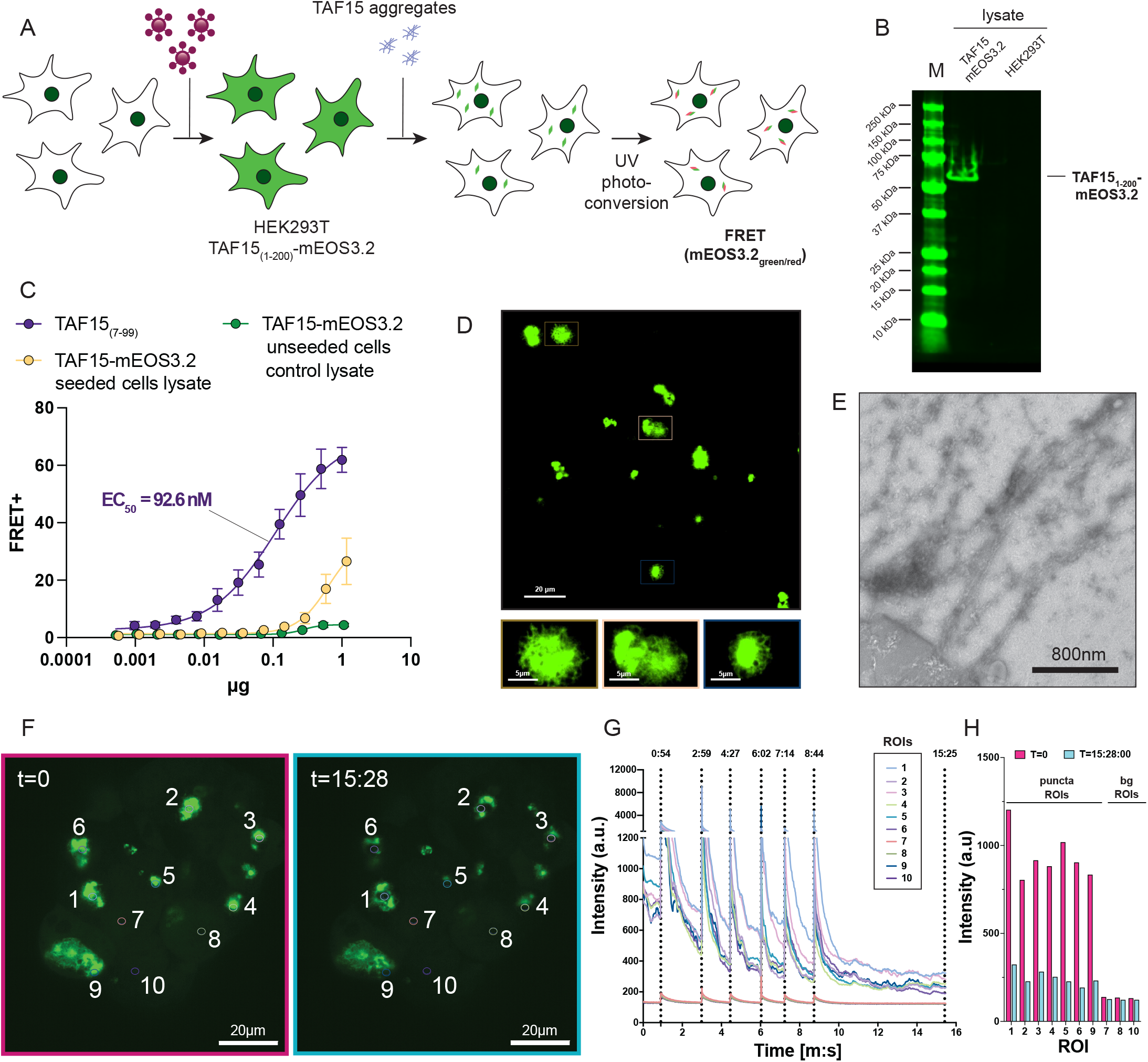
A single-fluorophore TAF15 biosensor reports cellular aggregation and serial propagation. **(A)** Design and workflow of the HEK293T biosensor expressing TAF15 LCD (residues 1–200) fused to mEOS3.2. **(B)** Fluorescent western blot of cell lysates showing the TAF15-mEOS3.2 band in biosensor cells (absent in HEK control lysate). **(C)** Flow-cytometry FRET readout of cells exposed to recombinant TAF15(7–99) seeds (shown in purple, n=4 individual replicates) and to “secondary” lysates harvested from previously seeded biosensors (shown in yellow, n=6 individual replicates), versus unseeded control lysate (shown in green, n=6 individual replicates). Dose–response to recombinant seeds yields EC_50_ ≈ 92 nM. **(D)** Confocal images of seeded biosensors showing cytoplasmic puncta with fibrillar substructure (scale bar: 20 μm); insets highlight fibril-like texture (scale bar: 5 μm). **(E)** TEM of sarkosyl-insoluble material extracted from seeded biosensors confirms intracellular amyloid fibril (scale bar: 800 nm). **(F–H)** FRAP of individual puncta: **(F)** pre- and post-bleach images with 10 ROI circles (scale bar: 20 μm); **(G)** fluorescence time courses per ROI **(H)** start/end intensity summary shows bleaching to background with no recovery, consistent with immobile fibrillar aggregates.

### Mapping TAF15 amyloid motifs

To identify the sequence determinants driving TAF15 aggregation and assess whether aggregation propensity is uniformly distributed across its amyloid core, we integrated predictive and experimental mapping. Utilizing the sequence-based algorithm WALTZ^48^ and the structure-based compatibility predictor CORDAX^49,50^, we identified several regions within residues 7-99 displaying high amyloid propensity (**Fig. 3A**). In particular, CORDAX predicted three segments spanning residues 16-21, 34-39 and 52-57. Overlapping predictions were highlighted by WALTZ for all three segments, together with an additional region encompassing residues 74-84. To complement these predictions, we applied thermodynamic profiling, using a previously established method^40,41,51^, to the patient-derived TAF15 amyloid fibril core structure (PDB ID: 8ONS). This method enables analysis of the per residue energetic contributions in fibril stability and has previously been shown to detect aggregation-prone regions (APRs) with very high accuracy^40,52^. Thermodynamic annotation identified four major stability hotspots within the amyloid core (**Fig. 3B**), largely overlapping with the computational predictions of overall aggregation propensity.

**Figure 3.**
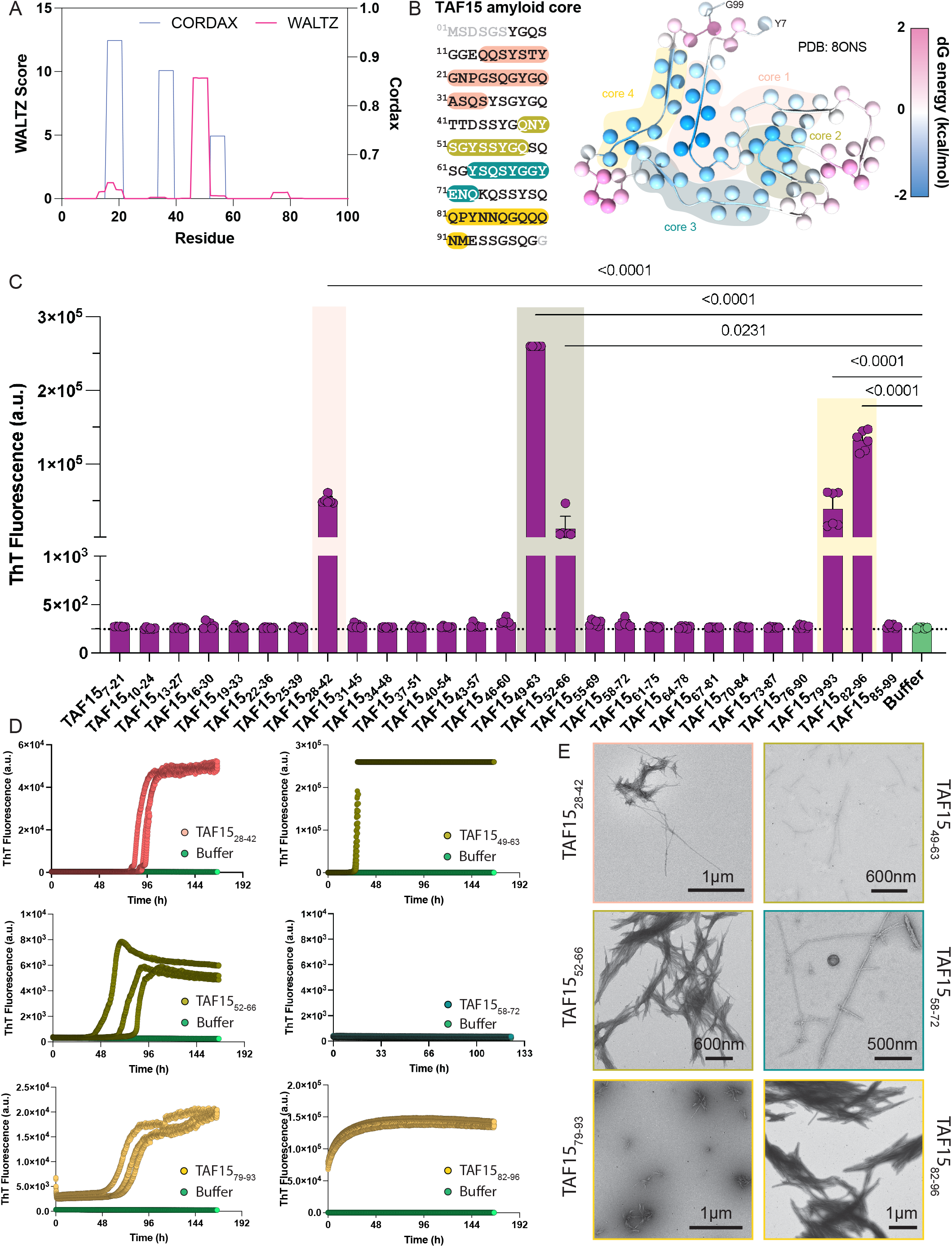
Integrated mapping of aggregation motifs in the TAF15 amyloid core. **(A)** Sequence-level amyloid propensity predicted across TAF15(1–100) from CORDAX (blue) and WALTZ (magenta), highlighting overlapping high-propensity segments. **(B)** Thermodynamic profiling of the patient-derived TAF15 fibril (PDB: 8ONS) maps per-residue stabilizing contributions; four stability hotspots are indicated and aligned to sequence. Endpoint ThT screening of a 27-member, 15-mer moving-window library (3-res step) spanning residues 7–99 identifies six ThT-positive windows; shaded boxes correspond to the hotspots in **(B)**. Statistics: One-way ANOVA with Dunnett’s multiple comparisons test (n=6 individual replicates). **(D)** ThT kinetics (n=3 individual replicates) of the six hit windows exhibit sigmoidal growth, color-coded to match panels **(A–C). (E)** TEM validates amyloid-like fibrils formed by each hit peptide; frame outlines retain the panel color code for cross-reference.

To experimentally determine the self-assembly propensity of these regions in an unbiased approach, we designed a moving-window peptide library spanning residues 7-99, composed of 27 overlapping 15-residue-long peptides, offset by a three-residue step. The aggregation propensity of each peptide was evaluated using ThT fluorescence assays. Our screen revealed five peptides, namely TAF15_28-42_, TAF15_49-63_, TAF15_52-66_, TAF15_79-93_, and TAF15_82-96_, displaying strong endpoint ThT signals, all mapping within the computationally predicted stability segments (**Fig. 3C**). Aggregation kinetics revealed characteristic sigmoidal ThT fluorescence curves for these peptides, but not for TAF15_58-72_, which corresponded to the fourth predicted stabilizing segment of the TAF15 fibril core (**Fig. 3D**). However, TEM imaging confirmed that all six peptides self-assemble leading to the formation of fibrils with canonical amyloid-like morphologies (**Fig. 3E**). Combined, our computational and experimental screened defined discrete APR motifs that correspond to the thermodynamic stable core regions of TAF15 amyloid fibrils.

### Self-propagation potential of TAF15 is encoded in amyloid motifs

Having defined six aggregation-prone motifs within the TAF15 amyloid core that overlap with thermodynamic stability hotspots, we next examined whether these segments could act as nucleation sites for assembly of the full TAF15(7–99) amyloid core fragment. In fluorescence ThT-based kinetic seeding assays, addition of preformed peptide aggregates markedly accelerated fibril formation of TAF15(7–99) in all cases (**Fig. 4A**). Endpoint fluorescence values confirmed that the active peptides efficiently templated amyloid formation, suggesting that apart of their role as energetically stabilizing regions, these sequences act also as self-assembly nucleators within the fibril core of TAF15.

**Figure 4.**
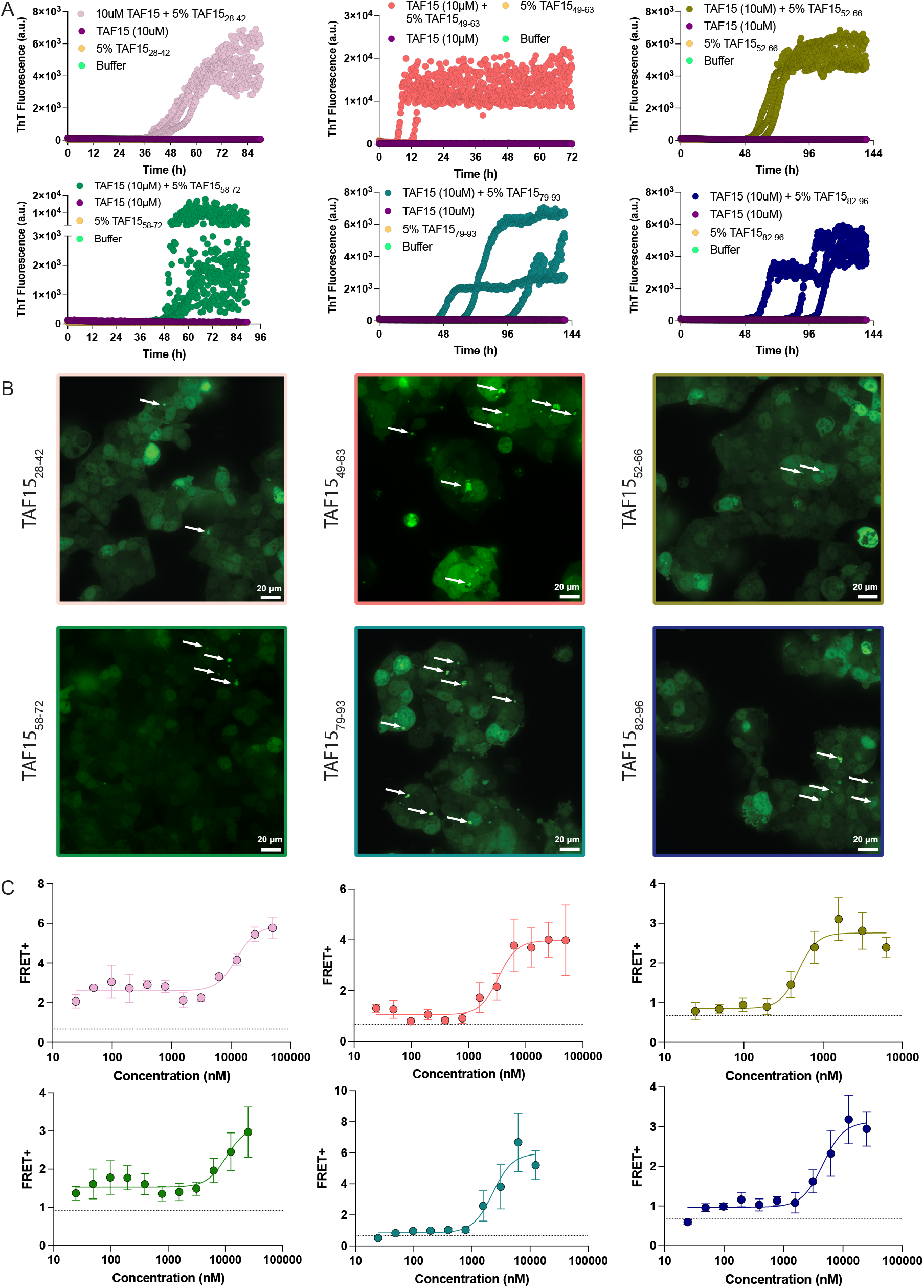
TAF15 motifs act as nucleation elements in vitro and in cells. **(A)** ThT seeding assays of TAF15(7–99) with aggregates from each APR peptide (n=3 individual replicates). All six windows accelerate aggregation relative to unseeded control. **(B)** Representative confocal imaging of TAF15 biosensor cells seeded with peptide-derived aggregates, yielding punctate cytoplasmic inclusions. **(C)** Quantification of cellular seeding (FRET) versus peptide-seed concentration reveals dose-dependent induction of cytoplasmic inclusions (n>4 individual replicates). Dotted line indicates control averages (untreated cells).

To assess whether these sequence motifs can also initiate aggregation in cells, we seeded the TAF15 biosensor with fibril preparations derived from each peptide. Confocal microscopy imaging confirmed the presence of punctate cytoplasmic inclusions with the fibrillar texture characteristic of seeded aggregates (**Fig. 4B**). Quantitative FRET analysis showed a concentration-dependent increase in signal for all peptides, consistent with what was observed in terms of their in vitro nucleating activity (**Fig. 4C**). Together, these results demonstrate that these identified aggregation-prone motifs are not only structural stabilizers within the amyloid core but further serve as functional nucleation elements capable of initiating templated TAF15 assembly. The strong concordance between energetic stabilization, in vitro seeding efficiency, and cellular propagation underscores that specific short sequence segments within TAF15 encode the molecular determinants of its amyloidogenic behavior.

### Specificity within the FET family and recruitment of FUS to TAF15 inclusions

To evaluate the selectivity of TAF15 seeding, we compared its aggregation behavior with that of other amyloidogenic proteins. Recombinant fibrils of tau(287-391), α-synuclein, and β-amyloid peptide (Aβ_42_) were generated and confirmed by TEM to display canonical amyloid fibril morphology (**Fig. 5A**). However, in contrast to TAF15 seeds, aggregates derived from these proteins failed to induce intracellular aggregation of TAF15(7–99), as reflected by the absence of a FRET signal in the TAF15 biosensor (**Fig. 5B**). These results confirm the high molecular specificity of the system for TAF15 assemblies.

**Figure 5.**
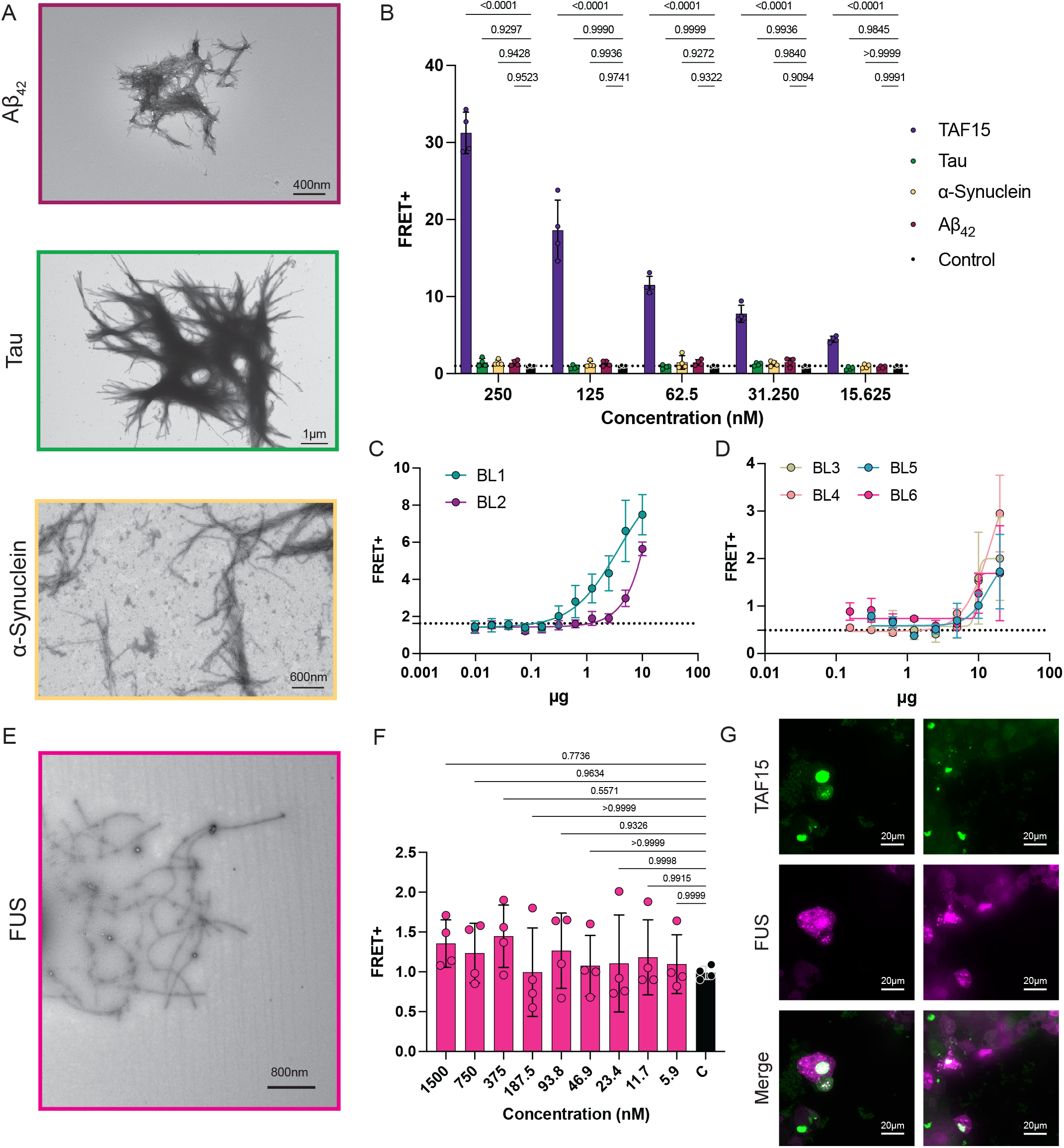
Specificity of TAF15 seeding and recruitment of FUS to TAF15 inclusions. **(A)** Electron micrographs of recombinant tau(287–391), α-synuclein, and Aβ_42_ fibrils used as heterologous amyloid controls. **(B)** TAF15 biosensor FRET readout shows no induction by heterologous amyloids (tau, α-synuclein, Aβ_42_) relative to TAF15 seeds, confirming molecular specificity. Statistics: Two-way ANOVA with t-tests for multiple comparison (n=4 individual replicates). Dotted line indicates control averages (untreated cells). **(C-D)** aFTLD-U brain lysates (n=4 individual replicates) seed the TAF15 biosensor in a concentration-dependent manner, demonstrating prion-like propagation of ex vivo TAF15 aggregates. Dotted line indicates control averages (untreated cells). **(E)** TEM validation of recombinant FUS fibrils displaying canonical amyloid morphology. **(F)** FRET quantification in the TAF15 biosensor demonstrating absence of cross-seeding by FUS aggregates (n=4 individual replicates). **(G)** Confocal micrographs of biosensor cells co-expressing mRuby-tagged FUS and seeded with TAF15 aggregates, showing co-localization of FUS within TAF15 inclusions.

To determine whether FTLD-FET patient-derived material exhibits similar prion-like propagation as observed for the recombinant TAF15 aggregates, we prepared lysates containing TAF15 aggregates from the prefrontal cortex of six individuals diagnosed with aFTLD-U (**Table S1**). These cases showed characteristic FUS pathology but stained negative for Alzheimer disease neuropathologic change (ADNC), TDP-43, and α-synuclein pathology. Lysates derived from all six individual aFTLD-U brains produced a clear, concentration-dependent increase in FRET signal in the TAF15 biosensor (**Fig. 5C-D**), demonstrating that pathological TAF15 aggregates from human FTLD-FET patient brains retain their prion-like cellular propagation capacity.

Given the close sequence homology and frequent pathological co-localization of TAF15 with FUS in FTLD– FET tissue, we next examined whether FUS assemblies could cross-seed TAF15 aggregation or be recruited to TAF15 inclusions. Despite the overall sequence similarity observed between TAF15 and FUS, recombinant FUS fibrils, produced and validated using TEM (**Fig. 5E**), did not nucleate TAF15 assembly in the biosensor cells (**Fig. 5F**). In contrast, expression of an mRuby-tagged FUS construct in the TAF15 biosensor, followed by seeding with TAF15 aggregates, showed partial co-localization of FUS within TAF15 inclusions (**Fig. 5G**). These observations indicate a largely unidirectional interaction between the two FET proteins: TAF15 aggregates do not depend on FUS for nucleation or cell propagation but can sporadically recruit FUS to growing inclusions. This behaviour aligns with neuropathological findings showing overlapping but not identical distribution of TAF15 and FUS immunoreactivity in pathological inclusions of FTLD–FET patients^6-9^, wherein amyloid filaments isolated from aFTLD-U brains are exclusively composed of TAF15^17^. Consistently, mass spectrometry analyses have shown no compositional differences in FUS peptides between affected and age control individuals, in contrast to TAF15^17^. Together, these results support that TAF15 acts as the core-forming species within FET inclusions, while FUS may be secondarily incorporated through heterotypic recruitment rather than through direct cross-seeding.

## Discussion

In this work, we establish TAF15 as a self-propagating amyloid capable of templated aggregation and prion-like propagation in cells. Using a combination of biophysical, computational, and cellular approaches, we define the minimal amyloid-competent region of TAF15, map its aggregation-prone sequence elements, and demonstrate that these motifs function as nucleation sites capable of driving both recombinant and cellular fibril formation. The mEOS3.2-based TAF15 biosensor developed here provides a quantitative platform to visualize and dissect the propagation behavior of TAF15 assemblies, enabling direct comparison with other amyloidogenic systems and patient-derived aggregates.

Thermodynamic profiling and peptide mapping revealed that the amyloidogenicity of the TAF15 low-complexity domain is encoded in a limited number of sequence hotspots that coincide with regions of enhanced structural stability in the aFTLD-U patient-derived TAF15 amyloid fibril fold^17^. This mirrors the organization of APR “hotspots” that stabilize protofilament cores in tau^32^, Aβ^53^, α-synuclein^54^, and TDP-43^55^, and extends the concept that amyloid cores are not a uniformly aggregation-prone entity but instead are composed of discrete sequence segments that nucleate and stabilize the pathological cross-β architecture^33,41^. The strong correlation between the computationally predicted, energetically stabilized, and experimentally validated segments underscores that amyloid-forming potential is a local property of the TAF15 sequence. Functionally, these motifs define the minimal energetic grammar that enables nucleation and propagation, suggesting that perturbations targeting these specific sites may provide a rational strategy to modulate TAF15 aggregation, in line with similar strategies undertaken for other key amyloid target proteins^43-45,56^.

At the cellular level, TAF15 aggregates exhibited self-templating behavior analogous to that of other well-characterized prion-like proteins, including tau and α-synuclein^57-59^, yet with a distinct molecular specificity. Patient-derived TAF15 aggregates seeded the TAF15 biosensor efficiently, whereas heterologous amyloids, including tau, α-synuclein, and Aβ_42_, failed to do so. This high selectivity reflects the strong sequence dependence of cross-β interface compatibility and may explain the restricted pathology spectrum associated with FET proteins. Despite the close sequence similarity between TAF15 and FUS, recombinant FUS fibrils were unable to seed TAF15 aggregation, while FUS recruitment to TAF15 inclusions was observed upon seeding. Such limited incompatibility within a single protein family contrasts with the partial cross-seeding observed among amyloid proteins sharing nucleating motifs with sequence similarity, as observed, for instance, for Aβ and medin^60,61^, as well as for other pathological amyloid proteins^43,62-64^, or even in cases of functional amyloid scaffolds^65-69^. This suggests that subtle differences in APR register or protofilament geometry could create a structural fence separating the two FET members. Indeed, alignment of the TAF15 and FUS low-complexity domains highlights that most of the short aggregation-prone motifs identified in TAF15 are not sequence-conserved in FUS, despite similarities in their enrichment of glycine, serine, and tyrosine residues. This divergence at the motif scale likely prevents formation of compatible steric-zipper interfaces, explaining the observed cross-assembly barrier. Nonetheless, multivalent LCD-LCD contacts could permit transient heterotypic interactions that recruit FUS to TAF15 inclusions, potentially modulating its assembly, as it has been observed for other amyloid forming proteins^60,61,70,71^. Together with neuropathological data showing co-localization of FUS and TAF15 but exclusive recovery of TAF15 filaments from patient tissue, these findings support a model in which TAF15 serves as the core-forming species within mixed FET inclusions, while FUS is incorporated through heterotypic recruitment.

More broadly, these results position TAF15 as a mechanistic link between the biophysical principles of low-complexity domain aggregation and the molecular selectivity of aggregates observed in FTLD-FET. By identifying the sequence determinants that govern TAF15 nucleation and propagation, this work provides a molecular framework to understand why TAF15, but not FUS or EWS, forms recoverable amyloid fibrils in disease. Notably, several ALS-associated TAF15 variants^22-29^, including A31T, N49S, S51G, E71G, and M92V, reside within or sit adjacent to the aggregation-prone segments defined in this work, suggesting that local energetic alterations in these regions could facilitate aberrant nucleation and accelerate alternative TAF15 pathology in ALS. Complementary studies have shown that TAF15 knockdown or parkin-mediated degradation mitigates TAF15-induced toxicity in ALS models^24^, reinforcing that aggregation of TAF15 itself is pathogenic and that modulation of its turnover may confer neuroprotection. These observations link the molecular determinants of aggregation defined here to potential therapeutic mechanisms and position TAF15 clearance pathways as promising targets. Future studies aimed at resolving the structural basis of TAF15–FUS interactions and defining how co-factors such as RNA might influence their co-assembly will further clarify how compositional diversity arises within FET inclusions and may suggest new points of therapeutic intervention.

## Materials and Methods

### Computational predictions and analysis

Calculations of aggregation propensity were performed using aggregation predictors WALTZ^48^ and CORDAX^49^. WALTZ employs a position-specific scoring matrix trained on a comprehensive set of amyloidogenic sequences^36^, while CORDAX is a structure-based predictor that captures amyloid propensity independently of beta-propensity and hydrophobicity^50^. Fibril stability was estimated using a recently developed thermodynamic profiling framework^33,40,41,52^, by summing all inter-residue interaction energies within the amyloid core of TAF15 fibril structures. Structural analysis and molecular graphics were generated using ChimeraX (v1.10.1), whereas plotting and statistical analyses were done using Graphpad Prism (v10.6.1).

### Protein expression, purification, and storage

A synthetic TAF15 (7–99) gene fragment (Twist Bioscience) was cloned into the pET-32a(+) vector and transformed into *E. coli* BL21(DE3) cells. Cultures (1 L LB + 100 µg/mL ampicillin) were grown at 37 °C to OD_600_ = 0.7–0.9, induced with 1 mM IPTG, and incubated overnight at 25 °C. Cells were harvested, resuspended in lysis buffer (50 mM Tris-HCl pH 8.0, 500 mM NaCl), treated with lysozyme (12 µg/mL, 30 min), sonicated, and clarified by centrifugation. The supernatant was applied to HisPur™ Ni-NTA resin (Thermo Fisher). The column was washed with lysis buffer containing 5 mM and 15 mM imidazole and eluted with 150 mM imidazole. The eluate was dialyzed overnight at 4 °C against Milli-Q H_2_O using Slide-A-Lyzer™ G3 (2 kDa MWCO) cassettes. The His-tag was removed by overnight digestion with His-tagged bovine enterokinase at room temperature. Cleaved protein was passed through Ni-NTA resin to remove the protease and TRX tag, concentrated using a 30 kDa MWCO filter, and lyophilized for storage (FreeZone Triad, Labconco). The tau fragment 287–391 was expressed in *E. coli* Rosetta (DE3) cells using LB auto-induction medium supplemented with 100 µg mL^−1^ ampicillin. Cultures were grown at 37 °C for 6–8 h, then at 24 °C for 18 h. Cells were harvested (5,000 × g, 15 min, 4 °C) and stored at –80 °C. Pellets were resuspended in lysis buffer (50 mM MES pH 6.0, 20 mM EDTA, 20 mM DTT, 0.2 mM PMSF, protease inhibitors) and lysed using a high-pressure homogenizer. Clarified lysate (14,000 × g, 40 min) was filtered and applied to a Capto S ion-exchange column pre-equilibrated in the same buffer. Proteins were eluted with a 0–1 M NaCl gradient and analyzed by SDS–PAGE. Pure fractions were pooled, further purified by size-exclusion chromatography (SEC), and stored at –80 °C in MES buffer. His-TEV-FUS-LCD (2–214) was expressed in E. coli BL21 (DE3) cells grown in LB medium containing 100 µg mL^−1^ ampicillin. Cultures were induced with 0.5 mM IPTG at OD_600_ ≈ 0.8–1.0 and incubated overnight at 20–25 °C. Cells were harvested (5,000 rpm, 20 min, 4 °C), washed with PBS, and stored at – 80 °C. Pellets were lysed in 50 mM Tris-HCl (pH 7.5), 500 mM NaCl, 1% Triton X-100, 20 mM β-mercaptoethanol, 2 M guanidine HCl, and protease inhibitors, with 0.2 mg mL^−1^ lysozyme. Lysates were sonicated, clarified (14,000 × g, 1 h, 4 °C), and purified using HisPur™ Ni-NTA resin (Thermo Fisher Scientific) with elution at 250 mM imidazole. The eluted protein was dialyzed against 20 mM Tris-HCl (pH 7.5), 200 mM NaCl, 20 mM β-mercaptoethanol, 0.5 mM EDTA, and 0.1 mM PMSF, then stored at –80 °C. Recombinant full-length α-synuclein was expressed and purified using a previously established protocol ^72^. Briefly, the E. coli BL21 (DE3) cells were transformed using the Alpha synuclein cloned pET28a plasmid, and the expression was induced using isopropyl-β-D-thiogalctoside (IPTG) (1mM). Cells were harvested, resuspended in lysis buffer (50mM Tris, 10mM EDTA, 150mM NaCl) containing protease inhibitor cocktail (Roche) and sonicated. The lysate was boiled at 95 °C for 20 min and centrifuged (7,000 × g, 30 min, 4 °C). The supernatant was treated with 10% streptomycin sulfate and glacial acetic acid, followed by ammonium sulfate precipitation (1:1 v/v). The complete precipitation was carried out by incubating for 4hr at 4°C. Precipitated protein was centrifuged and the pellet was washed with ammonium sulfate solution (1:1 (v/v) saturated ammonium sulfate and miliQ water). The washed pellet redissolved in ammonium acetate (100mM) and precipitated using ethanol. This step repeated twice followed by final resuspension in a small volume of ammonium acetate (100 mM) and lyophilization.

### SDS-PAGE

Samples (30 µL) from each purification step were mixed with 4× Laemmli buffer (Bio-Rad) containing 2-mercaptoethanol, heated at 95 °C for 5 min, and resolved on NuPAGE™ 4–12% Bis-Tris gels (Thermo Fisher). Precision Plus Protein™ All Blue Standards (Bio-Rad) were used as markers. Gels were stained with SimplyBlue™ SafeStain for 1 h, destained in H_2_O, and imaged on a Syngene G:BOX system.

### Western Blot

Total protein lysates (60 µg) from TAF15_1-200_–mEOS3.2 biosensor or HEK293T cells were mixed with 4× Laemmli sample buffer (Bio-Rad) containing 2-mercaptoethanol, heated at 95 °C for 5 min, and resolved on NuPAGE™ 4–12% Bis-Tris gels (Thermo Fisher Scientific). Proteins were transferred to nitrocellulose membranes (Trans-Blot Turbo Mini 0.2 µm; Bio-Rad) using the Trans-Blot Turbo transfer system. Membranes were blocked with 1% BSA in PBST and incubated with primary TAF15 antibody (1:1000; PA5-40335, Invitrogen), followed by Goat anti-Rabbit IgG (H+L) Alexa Fluor™ Plus 647 secondary antibody (1:5000; A32733, Invitrogen). Blots were imaged using an Azure Imager system (Azure Biosystems).

### Amyloid Fibril Formation and Fluorescence Assays

TAF15 (7–99) was dissolved in PBS (>1 mg/mL) and incubated at 37 °C with shaking (1,000 rpm) for 3–5 days. Peptides were diluted to 300 µM in 50 mM Tris (pH 7.4), 150 mM NaCl (7% DMSO). For fluorescence aggregation assays, samples were mixed with 25 µM ThT in 96-well plates (Greiner #675096) and incubated at 37 °C in a CLARIOstar plate reader (BMG Labtech). Endpoint fluorescence of TAF15 suspensions was measured after 5 days of incubation. For seeded reactions, 10 µM TAF15 was mixed with 5% (v/v) pre-formed fibril seeds and monitored as above. Tau fibrillization reactions were prepared by diluting purified Tau protein to 500uM in 10 mM phosphate buffer (pH 7.4) containing 10 mM DTT and 200 mM MgCl_2_. A total volume of 50 µL per reaction was transferred to 384-well plates. Samples were incubated at 37 °C under orbital shaking at 200 rpm using an Omega plate reader. His-TEV-FUS-LCD was aggregated at 2.5 mg mL^−1^ in 20 mM Tris-HCl (pH 7.5), 200 mM NaCl, 20 mM βmercaptoethanol, 0.5 mM EDTA, and 0.1 mM PMSF for 3–5 days at 37 °C with orbital shaking at 600 rpm. Finally, α-synuclein fibrils were prepared by incubating 5mg/mL protein in PBS at 37 °C with constant agitation (1,000 rpm) in an orbital mixer (Eppendorf). Reaction was stopped after 5 days of incubation. For cell seeding and TEM analyses, peptides (1 mM in Milli-Q H_2_O) were aggregated for 3–5 days at 37 °C with shaking (1,000 rpm). Seeds of TAF15 and of peptide amyloid fibrils were produced by sonication for 20 min at 65% amplitude using alternating 30 s on/off cycles in a QSonica bath sonicator.

### Peptide synthesis and purification

Peptides (blocked ends, N-terminal acetylation, C-terminal amidation) corresponding to the amyloid fibril core of TAF15 were synthesized on a MultiPep 2 robot (CEM) using Fmoc solid-phase synthesis with N-acetylated/C-amidated termini. Purity (>80%) was confirmed by HPLC (Shimadzu Nexera) and LC–MS (Shimadzu LCMS-2050). Peptides were ether-precipitated and stored at –20 °C. Peptide samples were pretreated with 1,1,1,3,3,3-hexafluoro-2-propanol (HFIP, Sigma-Aldrich), dried under nitrogen flow, redissolved in dimethyl sulfoxide (DMSO, Merck), filtered (0.2 µm), and diluted in final buffer. Aβ_42_ peptide was obtained from rPeptide and aggregated at a concentration of 1 mg mL^−1^ in PBS by incubating at 37 °C with shaking at 600 rpm for 5 days.

### Negative Staining

Ten microliters of each sample were applied to glow-discharged copper grids (Formvar/Carbon 400-mesh; Electron Microscopy Sciences) for 3 min. Grids were washed with Milli-Q water and stained with 2% (w/v) aqueous uranyl acetate for 1 min. Samples were imaged using a JEOL 1400 Plus transmission electron microscope operated at 80 keV.

### Thermodynamic stability of TAF15 amyloid fibrils

Chemical depolymerization and thermodynamic stability of TAF15(7-99) fibrils was determined using urea-induced disassembly coupled to Flow-Induced Dispersion Analysis (FIDA) of the soluble fraction (FidaBio). Preformed TAF15 fibrils (10 µM monomer equivalent) were incubated for at least 16 h at 25 °C in Milli-Q containing increasing concentrations of urea (0–6 M). Following incubation, samples were analyzed by FIDA to quantify the residual monomer fraction. The analysis was performed using the method below (Table 1) with a 100cm length and 75µm inner diameter capillary. For each elution profile, the fraction of soluble monomer (f) was calculated as the ratio of the area under the curve (AUC) corresponding to the monomer peak divided by the total integrated AUC for that sample. This value reflects the proportion of protein that remains in the soluble state under each denaturant condition. Thermodynamic analysis was performed using an isodesmic polymerization model. The global fit of all data points was performed by nonlinear least-squares minimization using the curve_fit function in SciPy (v1.13).

**Table 1.**
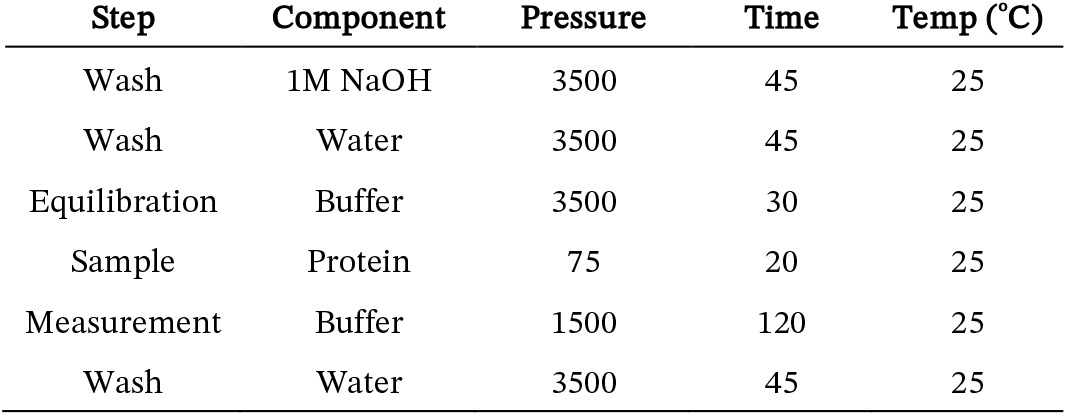
FIDA method for analysis of TAF15 amyloid fibril stability.

### TAF15_1-200_–mEOS3.2 Biosensor Cell Line Generation

A TAF15_1-200_–mEOS3.2 construct was cloned into a lentiviral transfer vector (Twist Bioscience). Viral particles were produced in HEK293T cells via triple transfection with psPAX2, pVSV-G, and the transfer plasmid using TransIT-293 reagent (Mirus Bio). Briefly, 3.5 × 10^5^ HEK293T cells were seeded in six-well plates and transfected the following day using TransIT-293 reagent (Mirus Bio) with 400 ng pVSV-G, 1,200 ng psPAX2, and 400 ng transfer plasmid per well. Viral supernatants were harvested 48 h post-transfection, clarified by centrifugation, aliquoted, and stored at –80 °C. HEK293T cells were transduced with lentivirus and cultured for 72 h. Single cells were isolated by FACS (FACSAria II, BD Biosciences). Clones were screened for responsiveness to TAF15 aggregates; the most sensitive clone was expanded and used for all subsequent assays.

### Cellular seeding assays

Proteins, peptides or lysates were mixed with Lipofectamine 2000 (0.5 µL per well) in Opti-MEM and reverse-transfected into 35,000 biosensor cells/well. After 48–72 h, cells were detached with Accumax, fixed in 1% PFA (PBS). Fixed cells were pelleted, resuspended in PBS + 3% FBS, and UV-illuminated for 40 min to photoconvert green mEOS3.2 to red. FRET was analyzed on an Attune NxT flow cytometer using excitation/emission 488 → 530/30 nm (green), 561 → 620/15 nm (red) and 488 → 695/40 nm (FRET). Data were processed in FlowJo. The gating strategy was as follows: HEK cells were first gated on FSC versus SSC to isolate the main cell population. From this population, single cells were selected using FSC-A versus FSC-H to exclude doublets and multiplets. Within the singlet gate, double-positive cells were identified based on green and red mEOS3.2 fluorescence. FRET-positive events were then defined as double-positive cells exhibiting increased signal in the FRET channel relative to negative controls.

### Neuropathological Analyses

Human brain tissue samples were obtained from three independent sources (Table S1): the University of Texas Southwestern Medical Center Neuropathology Brain Bank (UTSW), the Mayo Clinic Florida Brain Bank for Neurodegenerative Diseases (MCBB), and the Neurodegenerative Disease Brain Bank (NDBB) at the University of California San Francisco (UCSF).

At UTSW, each autopsy was performed with written permission of the decedent’s next of kin or another person legally authorized to provide consent according to Texas state law. All autopsies were restricted to brain examination only. In every case, one hemisphere was immersion-fixed in 20% neutral buffered formalin for at least 14 days after autopsy, while the contralateral hemisphere was flash-frozen and stored at –80 °C for future studies. Following fixation, each formalin-fixed brain specimen underwent comprehensive gross examination and dissection. Paraffin blocks were prepared for diagnostic evaluation, including sections from the hippocampus, middle frontal gyrus, superior and middle temporal gyri, inferior parietal lobule, calcarine cortex, cingulate gyrus, amygdala, lentiform nucleus/nucleus basalis of Meynert, thalamus/subthalamic nucleus, midbrain, pons, medulla, and cerebellum. All blocks were stained with hematoxylin and eosin for routine histopathological evaluation, and a standardized battery of histologic and immunohistochemical procedures was performed to characterize diagnostic lesions of Alzheimer’s disease (Alzheimer disease neuropathologic change, ADNC), non-AD tauopathies, synucleinopathies, TDP-43 pathology, FUS pathology, and vascular neuropathology. Diagnostic classification followed the current National Institute on Aging–Alzheimer’s Association (NIA-AA) guidelines^73^. All selected cases displayed neuropathologic features consistent with aFTLD-U.

For the UCSF cohort, postmortem brain tissue samples were collected and processed by the UCSF Neurodegenerative Disease Brain Bank (NDBB). Consent for brain donation was obtained from subjects or their surrogate decision makers in accordance with the Declaration of Helsinki and protocols approved by the UCSF Committee on Human Research. Clinical and neuropathological diagnoses were made as previously described^74^. Subjects were selected based on both clinical and neuropathological assessments and included individuals with a primary clinical diagnosis of behavioral variant frontotemporal dementia (bvFTD) without amyotrophic lateral sclerosis (ALS) and a neuropathological diagnosis of frontotemporal lobar degeneration (FTLD-FET) with atypical frontotemporal lobar degeneration with ubiquitinated inclusions (aFTLD-U). Demographic, clinical, and neuropathological information were included for all cases (**Table S1**).

At Mayo Clinic, all procedures involving human brain tissue were conducted in accordance with the ethical standards of the institutional and/or national research committees and with the 1964 Helsinki Declaration and its later amendments. Autopsies were performed with consent from the legal next of kin or another authorized representative. The collection and use of brain samples in this study were approved by the Mayo Clinic Institutional Review Board. Brains collected through the MCBB underwent systematic dissection and sampling following the established Alzheimer disease brain sampling protocol^75^. All individuals selected for this study exhibited neuronal cytoplasmic inclusions and occasional vermiform neuronal intranuclear inclusions in the dentate gyrus and frontal cortex that were immunoreactive for FUS (11570-1-AP, 1:500, rabbit polyclonal, Proteintech Group) and TAF15 (A300-308A, 1:500, rabbit polyclonal, Bethyl Laboratories). Haematoxylin and eosin staining did not reveal large basophilic inclusions or glassy hyaline conglomerate-like inclusions, and α-internexin immunohistochemistry (MCA-2E3, 1:100, mouse monoclonal, EnCor Biotechnology) did not demonstrate abundant neuronal intermediate filament inclusions. Based on these criteria, all selected cases displayed neuropathologic features consistent with atypical frontotemporal lobar degeneration with ubiquitinated inclusions (aFTLD-U).

### Brain Lysate Preparation

Brain tissue was homogenized in 1× TBS containing cOmplete™ protease inhibitor cocktail (Roche) at 10% (w/v) using a Power Gen 125 tissue homogenizer. Homogenates were sonicated for 5 min (amplitude 65%; 30s on/off), centrifuged (15,000 × g, 20 min, 4 °C). The supernatant was collected, and pellets were resuspended in 1 mL TBS + protease inhibitors, homogenized, and re-sonicated for 5 min. After a second centrifugation, both supernatants were combined, quantified (Pierce 660 Assay), aliquoted, and stored at –80 °C.

### Cell Lysate Preparation

For cell lysate seeding, 10^6^ biosensor cells were plated per well in a six-well plate and seeded with 1 µM TAF15_7-99_ fibril seeds for 48h. Cells were then homogenized in TBS + protease inhibitors, sonicated, centrifuged, and supernatants collected and stored as above.

### FUS expression and seeding

The FUS-LCD (1–214) gene block was synthesized by Integrated DNA Technologies (IDT) and cloned into an FM5-UBC-cassette-linker-Ruby vector containing a 12–amino acid flexible linker (GSAGSAAGSGEF). The construct was verified by Sanger sequencing. For transfection, 100 ng of plasmid DNA per well was reverse transfected into TAF15_1-200_–mEOS3.2 cells using 0.5 µL Lipofectamine 2000 (Thermo Fisher Scientific) per well. After 24 h, cells were seeded with 1 µM preformed TAF15 fibril seeds. Following a 48h incubation, cells were fixed with 4% paraformaldehyde (PFA) and imaged.

### Fluorescent Imaging and FRAP

Cell imaging of seeded biosensors was performed on a Nikon CSU-W1 SoRa spinning-disk confocal microscope (excitation 488 nm; emission 545 nm) at 37 °C with CO_2_ control. Using the same microscope, FRAP experiments were performed using a 100× oil immersion Plan Apo λ objective. Fluorescence recovery was recorded continuously for 929 s and laser power was set at 50%(Fig. S1). Live-cell imaging was carried out at 37 °C under controlled CO_2_ conditions. Peptide and FUS expression imaging was performed using an ImageXpress microscope equipped with a 60× Water Plan Apo VC objective. mEOS3.2-green fluorescence was acquired using excitation at 470 nm and emission at 520 nm, while FUS-LCD-Ruby fluorescence was imaged using excitation at 555 nm and emission at 624 nm.

### Extraction of TAF15 Biosensor Fibril Aggregates

Cell pellets were resuspended in filter-sterilized sucrose buffer containing 0.8 M NaCl, 10% (w/v) sucrose, 10 mM Tris-HCl (pH 7.4), 5 mM EDTA, and 1 mM EGTA. The suspension was homogenized using a Power Gen 125 tissue homogenizer and centrifuged (500 × g, 10 min, 4 °C). The supernatant was supplemented with 2% N-lauroylsarcosine (Sarkosyl) and incubated with glass beads at room temperature for 1 h on a nutator. Samples were centrifuged (186,000 × g, 60 min, 4 °C). The resulting supernatant was discarded, and the pellet was resuspended in 1 mL purification buffer containing 20 mM Tris-HCl and 100 mM NaCl.

## Acknowledgements

NNL and the Louros lab were supported by a scholarship from the Thomas O. Hicks Scholar in Medical Research. KK was partially supported by the O’Donnell Brain Institute (OBI) Sprouts grant program. Computational resources were provided by the BioHPC cluster supported by the Lyda Hill Department of Bioinformatics at UTSW. For cell sorting, we acknowledge the Moody Foundation Flow Cytometry Facility. We also acknowledge the assistance of the UT Southwestern Electron Microscopy Core, funded by the NIH grants 1S10OD021685-01A1 and to the UTSW Quantitative Light Microscopy Core for use of the Nikon SoRa spinning disk microscope, funded by NIH 1S10OD028630-01 to Dr. Kate Luby-Phelps. The authors want to thank Dr. Masato Kato and Dr. Steven McKnight for kindly sharing the His-TEV-FUS_(2-214)_ expression plasmid.

## Author contributions

KK and NNL conceived and initiated the study. KK, LG, and FG performed the experiments. NNL performed the computational analysis. KK and NNL performed data analysis. AG contributed to protein purification. KK and HDT contributed to peptide synthesis and purification. JVA and MD assisted with flow cytometry analysis. CLW, NBG, SFR, ALN, LTG, MAD, SES, WMS, and DWD provided critical patient tissue samples. NNL acquired funding and supervised the project. KK and NNL prepared the original draft and all authors contributed to the review and editing of the final version of this manuscript.

## Data Availability statement

All experimental and computational raw data generated, as well as un-cropped western blot membranes and SDS-PAGE gels are available as Source Data.

