## Supplementary Information for "TAF15 amyloids propagate via defined motifs in a prion-like fashion"

This PDF file includes:

Supplementary Table 1

Supplementary Figure 1

**Supplementary Table 1**. Clinical information of the patients and disease stage. An informed consent for autopsy and scientific use of autopsy tissue with clinical information was granted from all subjects involved.

| Sample ID | Source | Age | Sex | Neuropath Diagnosis/Disease Subtype | Clinical diagnosis | PMI (h) | Brain area |
|---|---|---|---|---|---|---|---|
| B1 | UTSW | 52 | M | FTLD-FET, aFTLD-U | Pick Disease | 10.5 | Temporal lobe |
| B2 | UTSW | 60 | F | FTLD-FET, aFTLD-U | Primary Progressive Aphasia | 5.6 | Temporal lobe |
| B3 | UCSF | 35 | M | FTLD-FET, aFTLD-U | bvFTD | 12 | orbitofrontal cortex, lateral |
| B4 | UCSF | 63 | F | FTLD-FET, aFTLD-U | bvFTD | 10.1 | orbitofrontal cortex, lateral |
| B5 | Mayo | 57 | F | FTLD-FET, aFTLD-U | CBS | 11 | Frontal cortex |
| B6 | Mayo | 48 | M | FTLD-FET/striatal atrophy/FBT/PNLA, aFTLD-U | bvFTD/PPA-NOS | 5 | Frontal cortex |

**Supplementary Figure 1.** Fluorescence recovery after photobleaching of TAF15 cellular inclusions. Individual panels indicate different timepoints of bleaching steps.

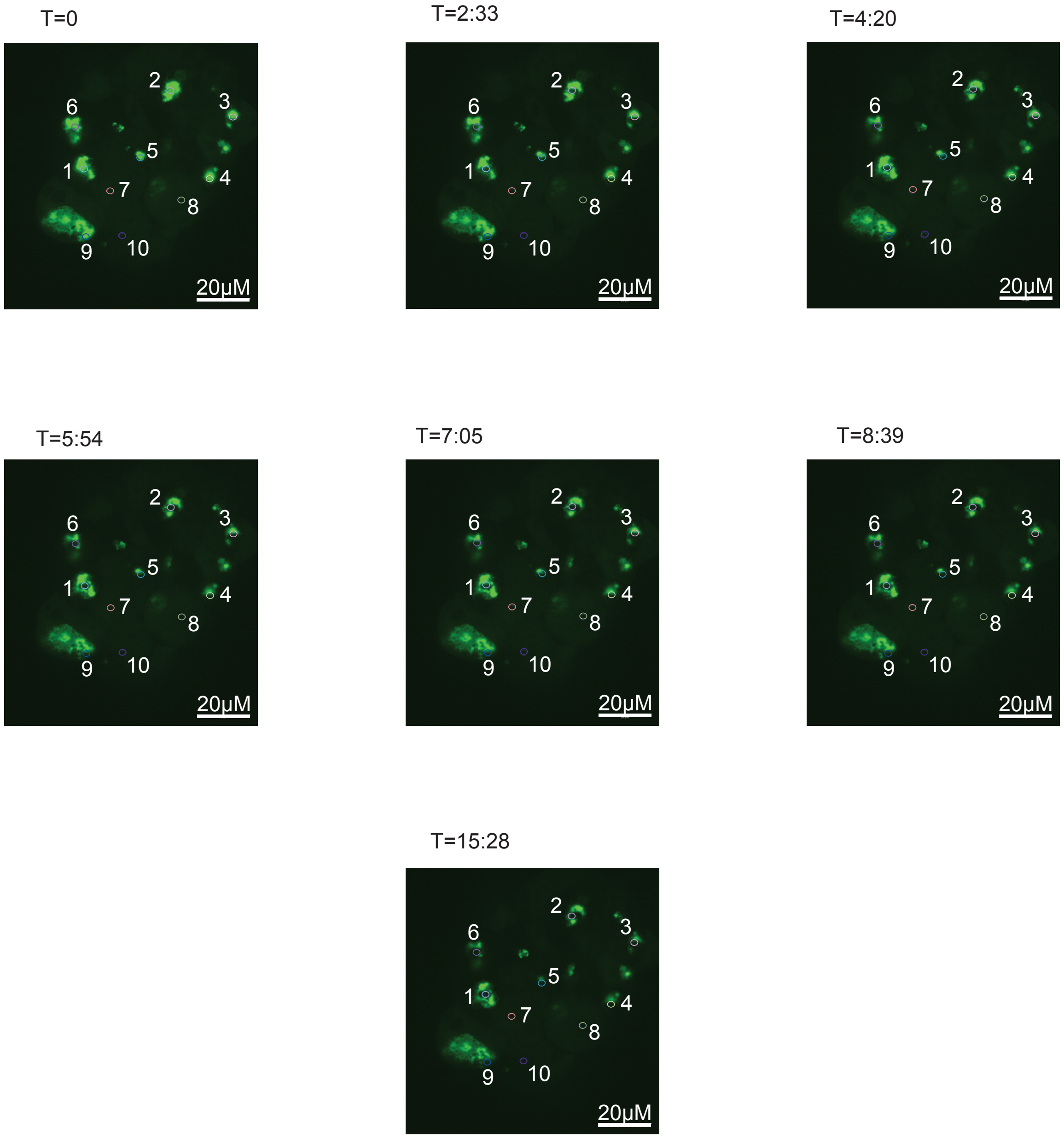
